# Towards prospective in-silico trials in atrial fibrillation: the case of polypharmacological SK and K2P channel block

**DOI:** 10.1101/2024.03.30.586087

**Authors:** Albert Dasí, Lucas M Berg, Hector Martinez-Navarro, Alfonso Bueno-Orovio, Blanca Rodriguez

## Abstract

**Background:** Virtual evaluation of medical therapy through human-based modelling and simulation can accelerate and augment clinical investigations. Treatment of the most common cardiac arrhythmia, atrial fibrillation (AF), requires novel approaches.

**Objectives:** To prospectively evaluate and mechanistically explain novel pharmacological therapies for atrial fibrillation through in-silico trials, considering single and combined SK and K_2_P channel block.

**Methods:** A large cohort of 1000 virtual patients was developed for simulations of AF and pharmacological action. Extensive calibration and validation with experimental and clinical data support their credibility.

**Results:** Sustained AF was observed in 654 (65%) virtual patients. In this cohort, cardioversion efficacy increased to 82% (534 of 654) through combined SK+K_2_P channel block, from 33% (213 of 654) and 43% (278 of 654) for single SK and K_2_P blocks, respectively. Drug-induced prolongation of tissue refractoriness, dependent on the virtual patient’s ionic current profile, explained cardioversion efficacy (atrial refractory period increase: 133.0±48.4 ms for combined vs. 45.2±43.0 and 71.0±55.3 for single SK and K_2_P block, respectively). Virtual patients cardioverted by SK channel block presented lower K_2_P densities, while lower SK densities favoured the success of K_2_P channel inhibition. Both ionic currents had a crucial role on atrial repolarization, and thus, a synergism resulted from the polypharmacological approach. All three strategies, including the multi-channel block, preserved atrial electrophysiological function (i.e., conduction velocity and calcium transient dynamics) and thus, its contractile properties (safety).

**Conclusion:** In-silico trials identify key factors determining efficacy of single vs combined SK+K_2_P channel block as effective and safe strategies for AF management.

## Introduction

Widely accepted in engineering, in-silico clinical trials now represent a new paradigm in medicine^1^. Similar to clinical trials, in-silico trials enable the assessment of medical interventions in a patient population, in this case composed of virtual patients using multiscale modelling and simulation. A regulatory framework for in-silico trials has been proposed^1^, and a growing body of literature is demonstrating their credibility and usefulness^2,3^. In-silico trials offer a wide range of advantages to augment clinical trials as well as preclinical investigations^1^: simulations can be conducted faster and with a lower economic burden, they are not subjected to ethical constraints, numerous interventions can be applied to the same virtual patient, and there is perfect control over the parameters of interest (i.e., less impact of confounding variables).

In this sense, in-silico trials have proven to be a powerful aid for assessing atrial fibrillation (AF) treatment, the most common cardiac arrhythmia^4^. Indeed, a large body of literature has demonstrated their credibility through consistency with experimental and clinical recordings^4-7^. Our recent study^8^ evidenced the ability of in-silico trials in large patient populations to predict and compare retrospectively the efficacy of 12 AF therapies (both pharmacological and ablation strategies). In addition to supporting the credibility of our approach, the simulations also identified key patient characteristics determining treatment success, which could guide stratification to optimal therapy. Thus, a further step for in-silico trials would be predicting the efficacy of interventions yet to be tested in large cohorts of patients (i.e., prospectively) to further strengthen the trustworthiness of these technologies, and to help refine the design of future clinical studies.

Recently, two clinical trials have been launched for two atrial-selective targets: the small-conductance Ca^2+^-activated K^+^ (SK) and the two-pore domain K^+^ (K_2_P) channels. Besides being predominantly expressed in the human atria, both the SK current (I_SK_) and TASK-1, a member of the K_2_P current (I_K2P_), are up-regulated in AF patients^9,10^. Accordingly, randomized clinical trials were registered to evaluate the cardioversion efficacy of single I_SK_^11^ and I_K2P_ inhibition^12^. The trials were unfortunately stopped prematurely (such as Covid-19-related slow recruitment^11^), and the results either consider a small sample size^11^ or are yet unpublished^12^. Since the available data can only be used to generate hypotheses^11^, further clinical trials with a bigger sample size are needed to determine the real efficacy of either treatment.

Thus, the aim of this study was to conduct a large-scale in-silico trial in a population of 1000 virtual patients to (1) assess the AF cardioversion efficacy of three pharmacological interventions (single SK inhibition, K_2_P inhibition, and combined SK+K_2_P channel block), and (2) demonstrate the power of in-silico trials for understanding cardiac arrhythmia mechanisms and selecting appropriate therapies.

## Methods

### Study Population

The construction of a similar cohort is explained in detail elsewhere^8^ and briefly summarized here.

The 1000 virtual patients were developed based on human data, to cover the variability in anatomy, electrophysiology and tissue structure commonly encountered in clinical practice (**Figure 1**). One half of the cohort presented structurally-healthy atria, and was developed combining 50 ionic current profiles and 10 atrial anatomies. These 500 virtual patients were then duplicated into a version that considered low voltage areas (LVA, bipolar voltage lower than 0.5 mV in sinus rhythm).

**Figure 1.**
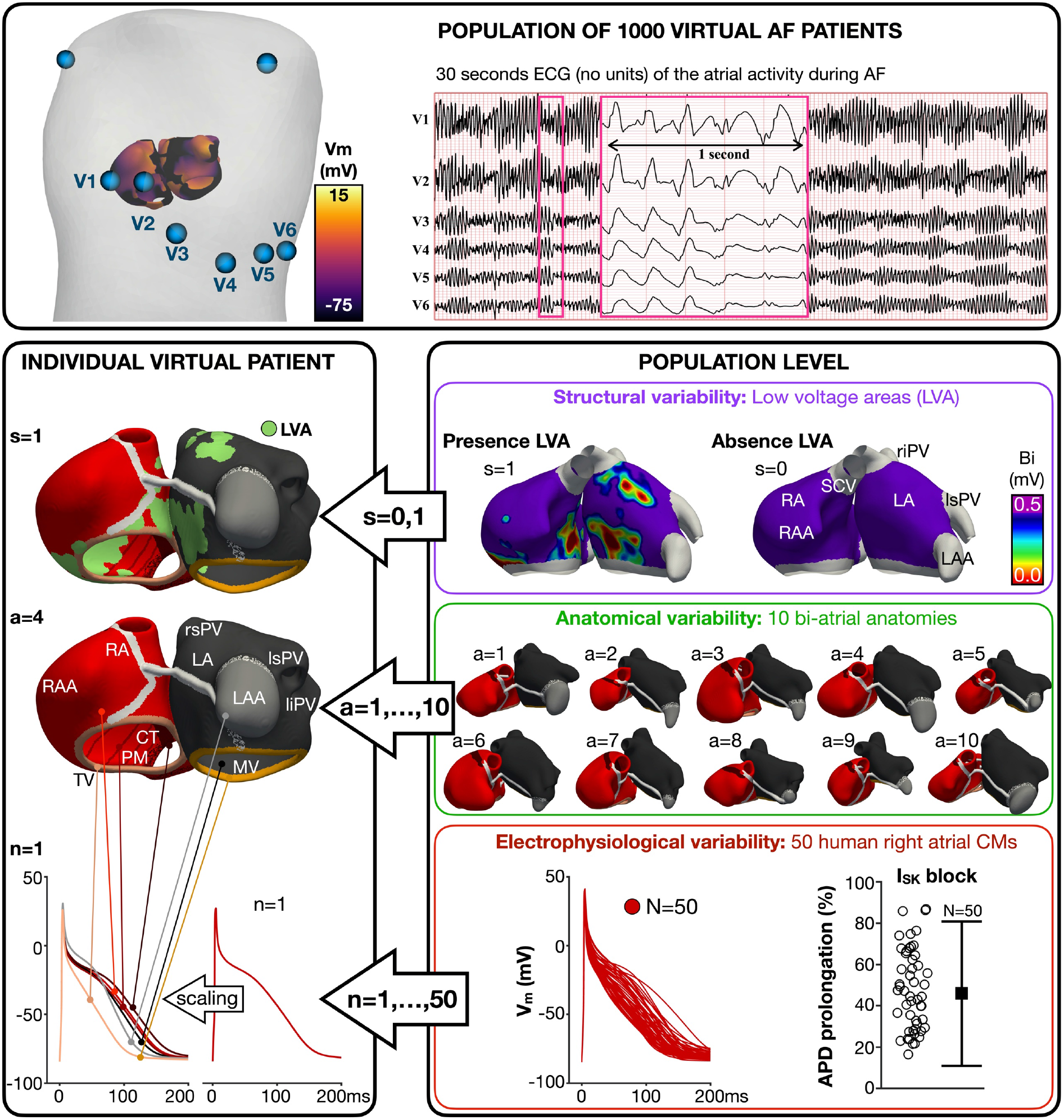
Construction of a population of 1000 virtual patients with atrial fibrillation (AF). **Top)** Representative virtual AF patient with the atria inside the torso. Screenshot of the simulated transmembrane voltage and 30 seconds ECG (dimensionless, only showing atrial activity). **Bottom-right)** Patients characteristics included at the population level: two different structural substrates, absence and presence of LVA; 10 different bi-atrial anatomies, with different right and left atrial volumes; and 50 different atrial cardiomyocyte models, with their corresponding action potential duration (APD) prolongation after SK channel block. **Bottom-left)** Construction of a unique virtual patient, including presence of LVA (s=1), the atrial anatomy number 4 (a=4) and the ionic current profile number 1 (n=1). The latter derives from taking the atrial cardiomyocyte model number 1 (n=1) and scaling it in six other atrial regions. **Abbreviations:** RA-LA: right and left atria; RAA-LAA: RA and LA appendage; TV-MV: tricuspid and mitral valve; CT: crista terminalis; PM: pectinate muscles; SCV: superior cava vein; rs-ri-ls-li-PV: right superior, right inferior, left superior and left inferior pulmonary vein.

#### Electrophysiological variability

A population of models approach^3^ was used to capture the ionic current variability commonly observed in experimental data. The population was constructed using a modified version of the Courtemanche et al.^13^ model, which included the K_2_P channel family, as in Wiedmann et al.^14^, and the SK channel, as in Celotto et al.^15^. Moreover, the maximal conductances of I_SK_ and I_K2P_ were scaled by 1.1 and 1.7, respectively, to reproduce the 12% and 18% action potential duration (APD) prolongation observed after single SK and K_2_P channel block in cardiomyocytes from control patients^9^.

Then, all ionic current densities of the modified model, including the SK and K_2_P channels, were systematically scaled in the range of [0.5, 2.5]. Five-hundred atrial cardiomyocyte models were developed following this process, each of them with a unique combination of ionic current densities. From this population, 50 cardiomyocyte models captured the variability in action potential characteristics of right atrial trabeculae from 149 chronic AF patients^16^ and the APD prolongation after SK channel block observed in right atrial cardiomyocytes from another six AF patients^9^ (**Figure 1, population level**). Each of these 50 models, which reproduced the single-cell behaviour of right atrial cells from two independent datasets of persistent AF patients, was further scaled to reflect electrophysiological heterogeneities in six atrial regions (**Figure 1, individual virtual patient**). The resulting seven action potential models (i.e., the original, representative of the right atrium, and the six scaled ones) configured a unique ionic current profile that was used to populate the atrial anatomies.

#### Anatomical variability

Ten atrial anatomies (with right and left atrial volumes of 127±51 mL and 105±39 mL, respectively; mean ± SD), spanning the volume ranges observed in AF patients, were selected. The 500 virtual patients with absence of LVA derived from the one-to-one combination of each atrial anatomy with each ionic profile.

#### Structural variability

These 500 virtual patients were duplicated into a version that included 15% LVA in the right and left atrium, as this LVA extension is associated with AF recurrence^8^. To incorporate LVA, the bi-atrial electro-anatomical maps of 20 (76% persistent) AF patients were used to build a probabilistic LVA map. The atrial regions most likely to undergo LVA remodelling were assigned to be LVA until 15% of the left and right atrium was remodelled. **Figure 1** shows the 15% bi-atrial LVA extension on a representative anatomy. LVA were simulated as regions of 30% decreased conductivity, increased anisotropy (i.e., 8:1 longitudinal to transversal conductivity ratio) and 50%, 40% and 50% reductions in I_CaL_, I_Na_ and I_K1_, respectively^4,8^.

### AF induction and ECG analysis

AF was induced in the population of 1000 virtual patients by imposing spiral wave re-entries as the initial conditions of the simulation^4,8^, since the main goal of the study was to assess AF cardioversion and not inducibility. AF lasting longer than 7 s was considered sustained, since we previously showed in a similar cohort^8^ that all AF episodes sustaining over 7 s would also sustain for 30 s (i.e., duration used clinically for AF diagnosis). Simulated 12-lead electrocardiograms (ECGs) were computed in virtual patients with sustained (>7 s) AF, and a representative example showing uninterrupted activity for 30 s is illustrated in **Figure 1**.

The three-dimensional monodomain equation of the transmembrane voltage and all ECG calculations were solved using the high-performance open-source MonoAlg3D^17^.

### Intervention

Virtual patients with sustained (>7 s) AF were independently subjected to three pharmacological interventions: complete inhibition (i.e., ionic current set to zero) of i) I_SK_, ii) I_K2P_, and iii) combined I_SK_+I_K2P_ block. The pharmacological interventions were modelled 2 s from AF initiation and the episode was continued until completion of the original 7 s (i.e., 2 s in the absence of intervention and 5 s after virtual drug administration^4^).

The primary outcome of the study investigated the proportion of virtual patients that were cardioverted before completing the original 7 s (i.e., cardioversion efficacy). The secondary outcome evaluated atrial cardiac safety. Safety regarding ventricular function and proarrhythmia (e.g., QT prolongation) has been already assessed in the clinical trials^11,12^. Thus, this investigation focused instead on assessing atrial function (calcium transient, conduction velocity, etc.) after I_SK_ and I_K2P_ block.

### Statistics

Data normality was assessed by the Kolmogorov–Smirnov test. Non-parametric data are presented as the median and interquartile range (IQR) and analysed using the Wilcoxon rank sum test. Parametric data are shown as the mean and standard deviation (SD) and analysed through the N-way analysis of variance. P<.05 was considered statistically significant.

## Results

### Characteristics of virtual patients with sustained AF

From the population of 1000 virtual patients, 654 (65%) presented sustained AF (> 7s). **Figure 2A** illustrates transmembrane voltage maps and corresponding atrial ECGs of two virtual patients, one with absence and one with presence of LVA. **Figure 2B** shows the number of AF episodes and rotors per episode according to the atrial volume for both subgroups of patients (absence and presence of LVA).

**Figure 2.**
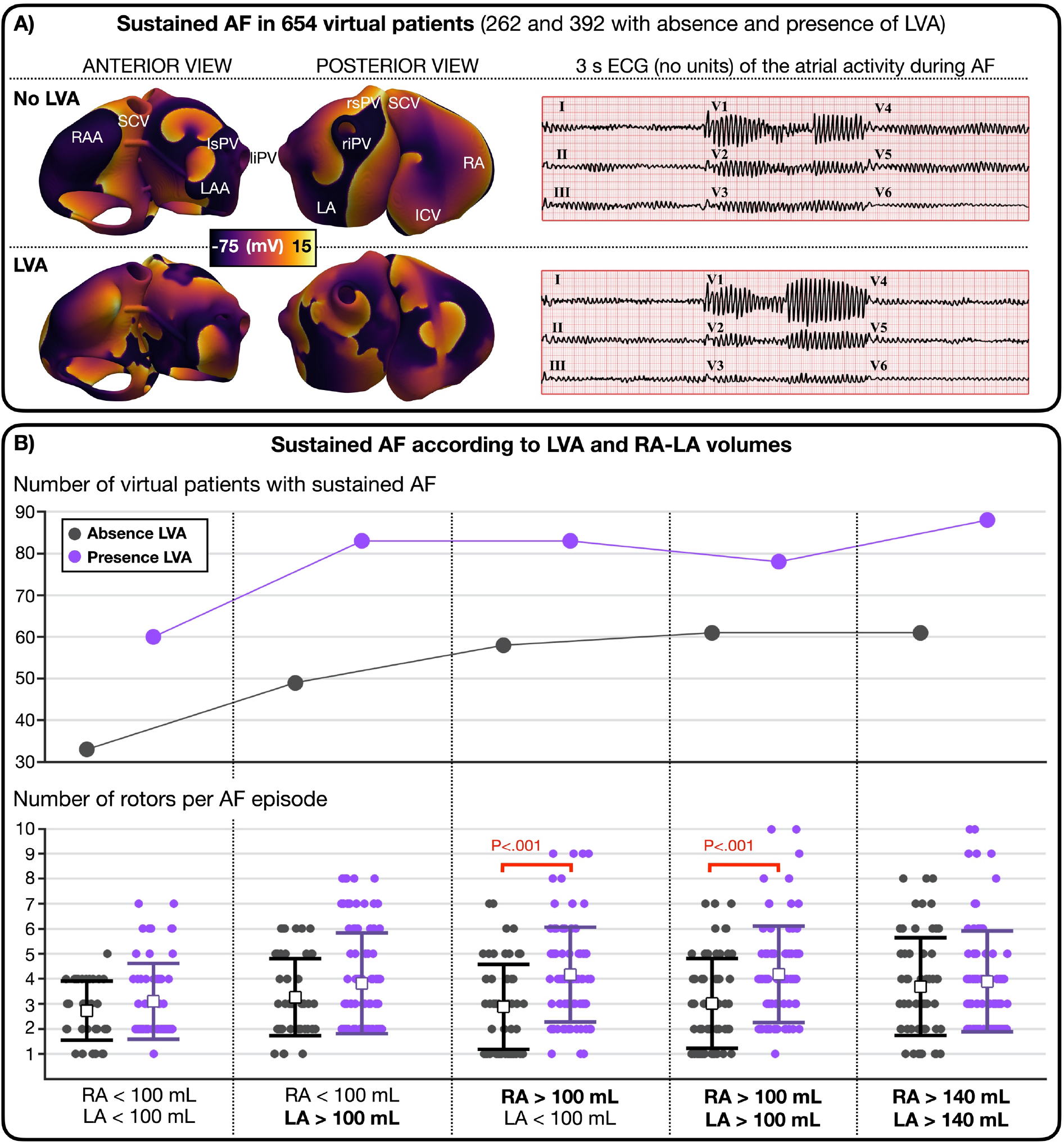
Comparison of atrial fibrillation (AF) dynamics between the subgroup of patients with absence and presence of low voltage areas (LVA). **A)** Screenshot of transmembrane voltage maps and electrocardiogram (ECG, only atrial activity) of a representative virtual patient with absence (no LVA) and presence of LVA. **B)** Number of virtual patients with sustained AF and number of rotors per AF episode according to increasing right (RA) and left atrial (LA) volumes (in mL).

AF maintenance was favoured by LVA and larger atria (**Figure 2B**). LVA led to more virtual patients sustaining AF (262 vs. 392 with absence vs. presence of LVA) and to an increase in AF complexity, qualitatively shown in **Figure 2A** and quantitatively evidenced by the increase in rotors per AF episode (3.1±1.7 vs. 3.9±1.9 with absence vs. presence of LVA; **Figure 2B**). Similarly, virtual patients with sustained AF were characterized by bigger atrial volumes (44% of virtual patients presented both atrial chambers above 100 mL, 42% at least one chamber over 100 mL, and only 14% had both chamber volumes below 100 mL, **Figure 2B**). Thus, more than half of the cohort with sustained AF (332 of 654) presented LVA and at least one atrial chamber enlarged.

### Efficacy endpoints

The 654 virtual AF patients were subjected to three independent pharmacological treatments, resulting in 2962 multi-scale simulations (i.e., 1000 in control conditions and 1962 after drug simulation). **Figure 3** illustrates the cardioversion efficacy obtained for the entire population (solid bars) and for both subgroups of virtual patients (dashed bars), with absence (262 of 654) and presence (392 of 654) of LVA.

**Figure 3.**
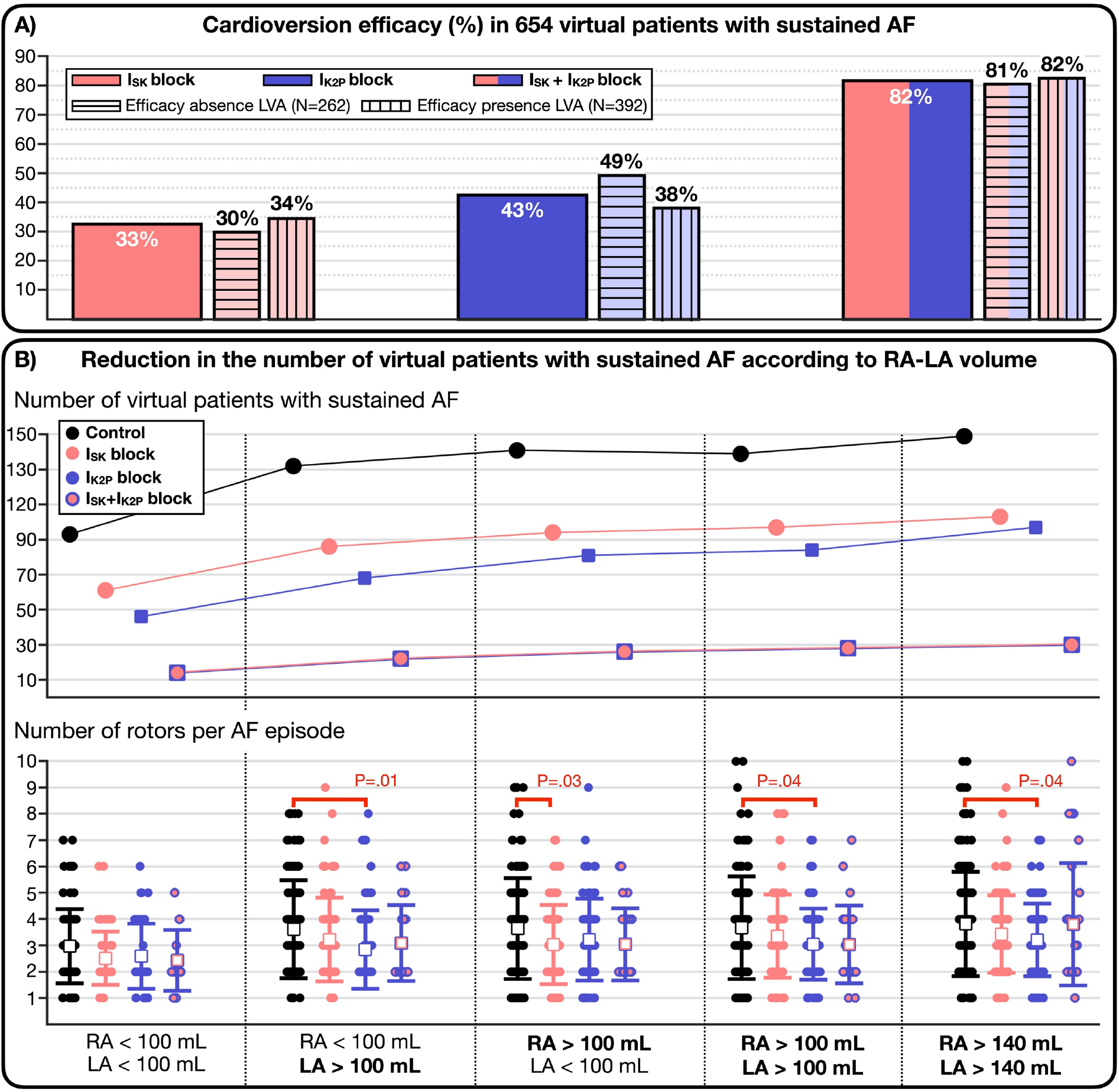
Cardioversion efficacy of single and combined SK and K_2_P channel block. **A)** Cardioversion efficacy in the 654 virtual patients with sustained atrial fibrillation (AF) (solid bars) and in both subgroups of virtual patients (dashed lines), with presence and absence of low voltage areas (LVA). **B)** Number of virtual patients with sustained AF in control conditions and after the application of the three virtual treatments according to increasing right (RA) and left atrial (LA) volumes (in mL). Number of rotors per AF episode in those virtual patients non-cardioverted by each treatment.

Single I_SK_ and I_K2P_ block exhibited limited cardioversion efficacy, terminating AF in 33% (213 of 654) and 43% (278 of 654) of virtual patients, respectively. Remarkably, combined I_SK_+I_K2P_ inhibition had a higher cardioversion efficacy than adding the success rate of single channel blocks (i.e., synergism), stopping AF in 534 (82%) cases.

A similar efficacy derived from single SK and K_2_P channel inhibition (33% vs. 43%, respectively). However, from the 213 virtual patients responding to I_SK_ block and the 278 ones responding to I_K2P_ block, only 108 virtual patients were cardioverted by both treatments. Conversely, 105 (49%) and 170 (61%) virtual patients cardioverted by single I_SK_ and I_K2P_ inhibition, respectively, did not respond to the other treatment. Neither strategy showed a strong dependency between cardioversion efficacy and atrial volume (i.e., a similar proportion of virtual patients with different atrial volumes were cardioverted, **Figure 3B**), and only the efficacy of I_K2P_ block was considerably higher in the absence of LVA (**Figure 3A**). Furthermore, both treatments, especially I_K2P_ block, significantly reduced the number of rotors per AF episode in those virtual patients that were not cardioverted (**Figure 3B**).

Compared to single channel block, combined I_SK_+I_K2P_ inhibition yielded a greater reduction in the number of virtual patients with sustained AF (**Figure 3B**). Indeed, besides cardioverting AF in the majority of cases where single channel block was successful (207 of 213 and 260 of 278 for I_SK_ block and I_K2P_ inhibition, respectively), the synergistic strategy also terminated AF in 174 additional virtual patients in which neither I_SK_ block nor I_K2P_ inhibition worked on isolation.

### Mechanisms of cardioversion

**Figure 4A** illustrates the drug-induced prolongation in APD and effective refractory period (ERP) for all three pharmacological strategies. Longer APDs were observed after combined I_SK_+I_K2P_ block, followed by single I_SK_ inhibition and I_K2P_ block. When the atrial cardiomyocyte models were paced at 1 Hz, all APD comparisons yielded statistical significance (not included in **Figure 4A**) and I_SK_ inhibition prolonged the APD to a greater extent than I_K2P_ block. For 4 Hz pacing, however, no significant difference was obtained between I_SK_ block and I_K2P_ inhibition, and the combined strategy led to a greater APD prolongation than single channel block (**Figure 4A**). All three pharmacological treatments significantly prolonged the APD compared to control conditions. At tissue level, the increase in the ERP was proportional to the cardioversion efficacy: higher for combined I_SK_+I_K2P_ block, followed by I_K2P_ inhibition and, lastly, I_SK_ block (**Figure 4A**).

**Figure 4.**
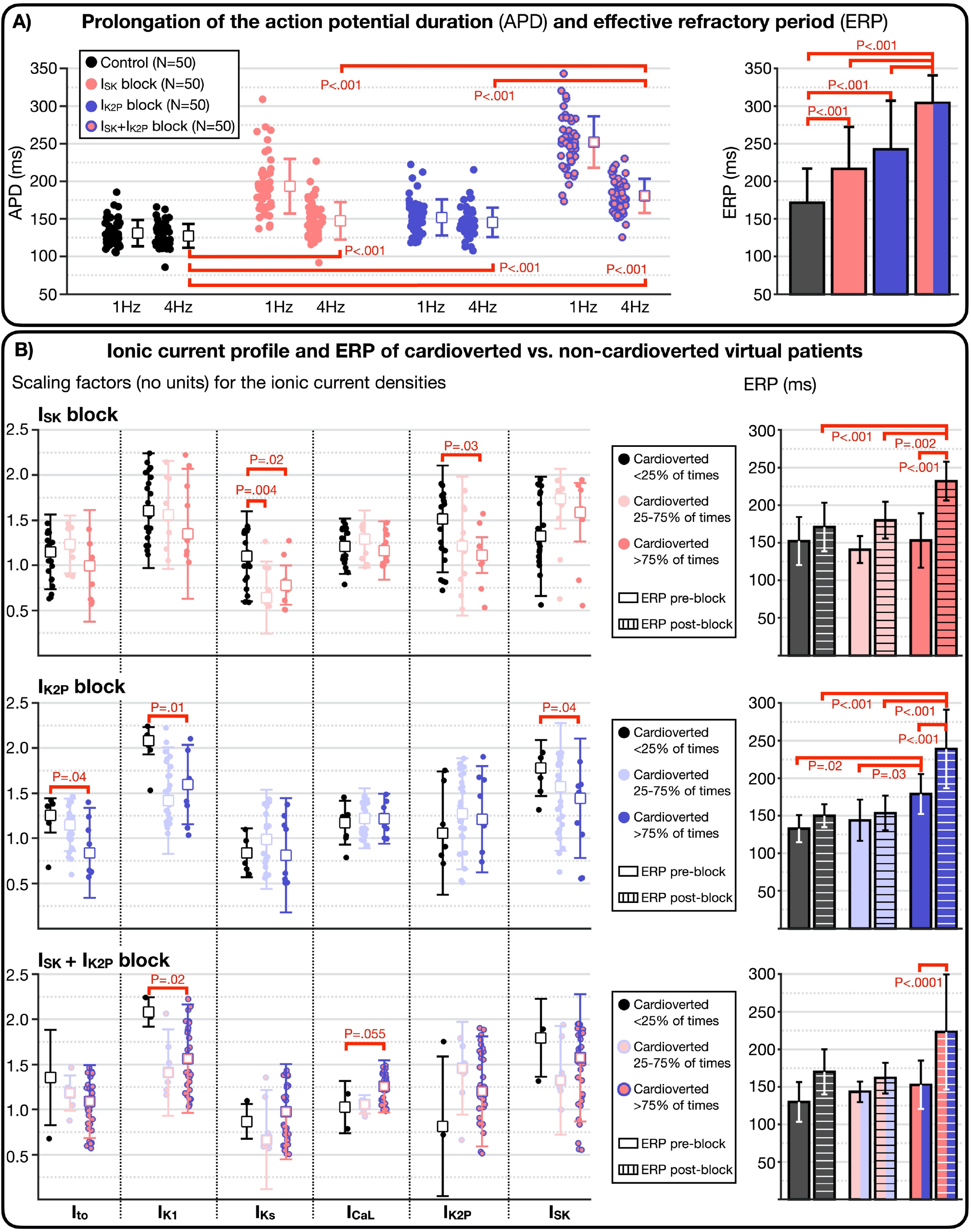
Mechanisms of atrial fibrillation (AF) cardioversion. **A)** APD, computed in single-cell at 1 and 4 Hz, and ERP, computed in tissue, of the 50 atrial cardiomyocyte models in control conditions and after applying all three pharmacological treatments. **B-left)** Scaling factors (dimensionless) for the ionic current densities used in ionic profiles that were cardioverted less than 25% of times, between 25% and 75%, and more than 75% of times after applying each pharmacological treatment. **B-right)** ERP of ionic current profiles that were cardioverted less than 25% of times, between 25% and 75%, and more than 75% after applying each pharmacological treatment. The ERPs are shown before (pre-block, solid bars) and after (post-block, dashed bars) applying each strategy.

**Figure 4B** compares the ionic current profile and the drug-induced ERP prolongation between cardioverted and non-cardioverted virtual patients. Of importance for the analysis, since the same ionic current profile can be used in 20 different virtual patients (i.e., combination of the ionic profile with each of the 10 atrial anatomies, with absence or presence of LVA), **Figure 4B** considers three pooled groupings: **i)** ionic current profiles rarely cardioverted by the drug (i.e., cardioversion occurred in less than 5 (25%) of the 20 virtual patients sharing the same ionic profile); **ii)** ionic current profiles usually cardioverted by the drug (i.e., cardioversion occurring in between 5 and 15 (25-75%) virtual patients with the same profile); and **iii)** ionic current profiles with frequent cardioversion (i.e., cardioversion in more than 15 (75%) virtual patients with the same ionic profile). In the text below, virtual patients with ionic current profiles that achieved more than 75% cardioversion are referred as cardioverted virtual patients, and the remaining patients are considered non-cardioverted.

Generally, for all three pharmacological strategies, virtual patients showing successful cardioversion also experienced a greater prolongation of refractoriness after the application of the virtual treatment (i.e., ERP post-block, **Figure 4B**). For single I_SK_ and I_K2P_ block, the ERP post-block of cardioverted patients was, not only significantly longer than their corresponding ERP pre-block but also, significantly longer than the ERP post-block of non-cardioverted virtual patients. Interestingly, virtual patients cardioverted by I_K2P_ block also presented a longer ERP pre-block than non-cardioverted virtual patients. This explains that blocking I_K2P_ was more efficacious in the absence of LVA (**Figure 3A**), in which the ERP was the main determinant of AF maintenance.

The most remarkable finding was that virtual patients frequently cardioverted by I_SK_ block presented lower I_K2P_ density (alternatively, high I_K2P_ hindered the success of I_SK_ block), and similarly, I_K2P_ inhibition was successful in virtual patients with lower I_SK_ density (alternatively, high I_SK_ hampered I_K2P_ block). This finding suggests that, since both ionic currents had an important role on tissue refractoriness, blocking one current was not effective when the other one was up-regulated. Moreover, it explains the high efficacy of the synergistic block, remaining only unsuccessful in virtual patients with shorter refractoriness (due to concomitant I_K1_ up-regulation and I_CaL_ down-regulation, see **Figure 4B**).

Additionally, I_SK_ block was less efficacious in virtual patients with I_Ks_ up-regulation and with a non-significant tendency for lower I_SK_. On the other hand, I_K2P_ inhibition had a lower efficacy in virtual patients with increased I_to_ and I_K1_ (**Figure 4B**). As expected, the cardioversion efficacy of single channel block was less efficacious in virtual patients with a strong repolarization reserve.

### Safety endpoints

Neither pharmacological treatment produced a detrimental change on the atrial electrophysiological function (**Figure 5**). No differences were observed in the calcium transient duration or in the longitudinal conduction velocity in the bulk tissue after I_SK_ and I_K2P_ inhibition compared to control conditions. Only the calcium transient amplitude and the diastolic level were slightly higher after SK channel block and thus, also after combined I_SK_+I_K2P_ inhibition (**Figure 5**).

**Figure 5.**
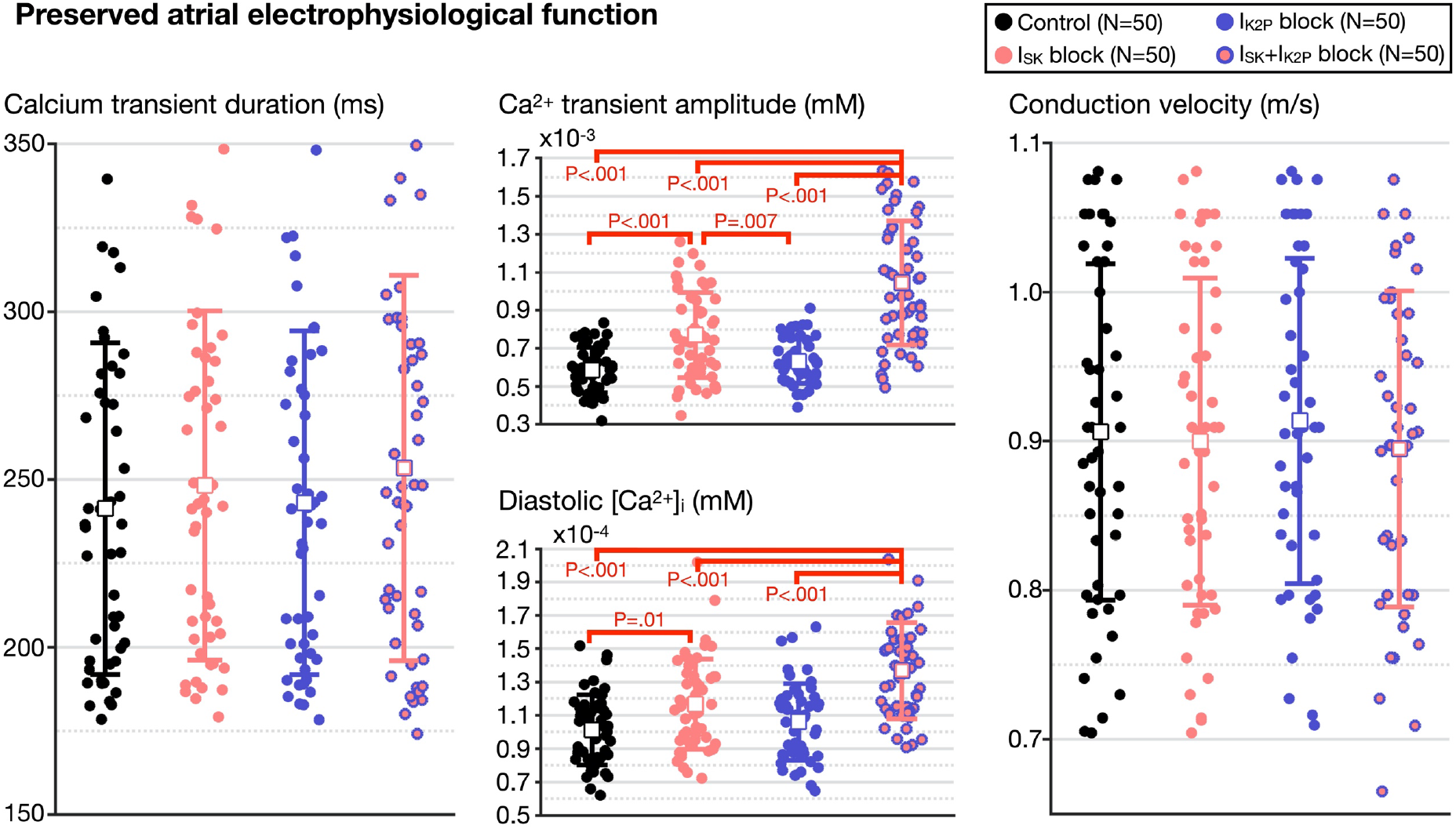
Safety endpoints of SK and K_2_P channel block. Comparison of calcium transient biomarkers and longitudinal conduction velocity in the bulk atrial tissue between control conditions and after the virtual application of single and combined SK and K_2_P channel inhibition.

## Discussion

This is a prospective, large-scale in-silico trial study considering 1000 virtual patients to evaluate single and combined application of two new therapies (SK and K_2_P channel inhibition) for AF. In-silico trials exploit the complete control and transparency of modelling and simulation to identify key factors determining response to different therapies in the same cohort of virtual patients. Our results reveal that AF cardioversion efficacy was proportional to an increase in tissue refractoriness, determined by the specific pharmacological therapy and the underlying patient’s ionic profile. When considering single channel block, I_K2P_ and I_SK_ up-regulation had a crucial role on the success of pharmacological treatment, which emphasised the benefits of adopting a polypharmacological strategy.

### Comparison to preclinical and clinical studies

We previously conducted in-silico trials in a similar population of 800 virtual patients to assess optimal pharmacological and catheter ablation therapy for AF^8^. The credibility of the in-silico trials was supported by consistency with experimental and clinical data, and also with the efficacy reported by 12 therapies and their corresponding clinical trials and studies^8^.

For the current study, a larger population of 1000 virtual patients was developed, to include the single-cell effects of SK channel block observed in atrial cardiomyocytes from persistent AF patients^9^. Compared to the 33% (213 of 654) of virtual patients cardioverted in-silico, AF cardioversion rates of 43% (5 of 12) and 55% (12 of 22) have been reported in a phase-2 clinical trial of AP30663 (SK channel blocker) at 3 and 5 mg/kg, respectively^11^. The higher cardioversion efficacy observed in human patients might be due to many reasons, for instance, the small sample size of the trial, the fact that only paroxysmal AF patients were enrolled, or that besides inhibiting the SK channel, AP30663 also produces an off-target inhibition of Kv11.1 (hERG gene)^11^.

Nonetheless, the in-silico I_SK_ block reproduced an ideal scenario, where a complete SK channel inhibition was achieved. In a study conducted in pigs^18^, 5 mg/kg of AP30663 produced a free plasma concentration equivalent to the IC_50_ of the SK channels (i.e., 50% inhibition of I_SK_). Thus, an even higher cardioversion efficacy could have been obtained in the clinical trial if a higher dose (i.e., 25 mg/kg) had been tested^11^. In pigs^18^, a dose of 25 mg/kg yielded a 71% (+83±7 ms) increase in the atrial ERP. This is in agreement with the ERP prolongation (+73.6±47.4 ms) seen in the current study for cardioverted virtual patients. To the best of our understanding, the atrial ERP modulation by SK channel block has not been reported in humans, but our in-silico cardiomyocyte models reproduced the APD prolongation observed in patients with AF^9^.

Regarding I_K2P_ inhibition, the results from the DOCTOS trial^12^ are yet to be published. In animal studies, a 100% cardioversion rate was observed in 17 pigs that received 1.8 mg/kg of doxapram (TASK-1 inhibitor)^19^. At a similar dose (2.0 mg/kg), the porcine atrial refractory period was prolonged on average by 38.3, 25.8 and 42 .5 ms at cycle lengths of 500, 400 and 300 ms, respectively^19^. In our simulations, a further ERP prolongation of 71.0±55.3 ms was observed in human virtual patients. In isolated cardiomyocytes from persistent AF patients, I_K2P_ inhibition increased the APD up to 30%^10^, compared to an average and a maximum prolongation of 16±8% and 34.4%, respectively, observed in our simulations. Thus, we anticipate that the DOCTOS trial will report a comparable efficacy in AF patients as that observed in-silico after K_2_P channel block.

### Comparison to previous simulation studies

A previous study by Ni et al.^20^ already hypothesized that the combined block of several atrial-selective K^+^ currents would provide a synergistic antiarrhythmic effect. The authors used cable tissues to assess the impact of single and multiple channel block on the APD, ERP and conduction velocity. We simulated AF in anatomical atrial meshes and, in agreement with Ni et al.^20^, a greater ERP prolongation was observed for the combined I_SK_+I_K2P_ block, followed by single inhibition of I_K2P_ and, lastly, I_SK_ block. Furthermore, they observed a negligible effect on the conduction velocity after blocking these channels, also shown in the current study (**Figure 5**).

Another and more recent study by the same group^21^ observed that under AF conditions, I_SK_, I_K2P_ and I_K1_ were the K^+^ currents with the most prominent role in the atrial ERP. Consistently, we observed that I_SK_ and I_K2P_ up-regulation dictated the success of pharmacological cardioversion, and when both ionic currents were inhibited, cardioversion only failed in those virtual patients with I_K1_ up-regulation. Furthermore, the authors observed that increased SK channel density reduced the amplitude of the calcium transient, or equivalently as we observed, increased calcium transient amplitude derived from I_SK_ block (**Figure 5**).

As in the present study, both works^20,21^ used a population of cardiomyocyte models approach to include electrophysiological variability. Importantly, the authors employed a different cellular model to build the population, which demonstrates model independence of key results.

### Efficacy and safety benefits of multi-targeting atrial-selective ionic currents

With the completion of preliminary trials assessing I_SK_ block^11^ and I_K2P_ inhibition^12^, all atrial-selective ionic currents, including the acetylcholine-dependent K^+^ current (I_K,ACh_)^22^ and the ultra-rapid rectifier K^+^ current (I_Kur_)^23^, have been evaluated for rhythm control of AF in randomized clinical trials.

I_Kur_ was considered a very attractive target since the inhibition of this major repolarization current in the human atria was expected to prolong the ERP and destabilize rotors. The DIAGRAF-IKUR trial^23^, however, ended prematurely, showing no meaningful reduction in disease burden in paroxysmal AF patients. Several reasons were given to explain the limited success of I_Kur_ block, for example, some studies demonstrated a decreased I_Kur_ density in cardiomyocytes from AF patients^24^.

This was not the case for I_K,ACh_ (also I_SK_ and I_K2P_), whose activity appeared consistently up-regulated in AF patients^24^. Yet, the trial assessing I_K,ACh_ inhibition also reported a lack of efficacy for reducing AF burden^22^. The authors speculated that the compound used might have not blocked I_K,ACh_ in AF patients. A more interesting reason, also highlighted by the investigators of the DIAGRAF-IKUR trial^23^, was that meaningful AF prevention needed more than the single inhibition of one atrial-selective current.

In agreement with the latest statement, we have demonstrated that targeting multiple atrial-selective ionic currents results in a synergistic cardioversion efficacy (i.e., greater than adding the individual efficacy of single channel block). Moreover, the benefits of multiple channel block could also be extended to cardiac safety. For example, ventricular cardiomyocytes from heart failure patients show I_SK_ up-regulation (extensively reviewed by Herrera et al^21^). Thus, I_SK_ inhibition might be associated with safety concerns in AF patients with concomitant heart failure. Moreover, both doxapram^19^ (I_K2P_ inhibitor) and AP30663 (SK channel blocker)^11^ produce an off-target effect on the hERG current at high doses. Thus, a lower-dose, multi-target strategy would be more efficacious and safer.

## Limitations

This study aimed to study AF cardioversion in a large population of virtual patients. Thus, a compromise was reached between the total number of simulations (i.e., 2,962) and the duration of each simulation (i.e., 7 seconds). However, it is unclear whether 7s of simulated AF is equivalent to the 90 minutes waiting reported in the trials^11^ and thus, if could be used to predict AF cardioversion. In this sense, since pharmacokinetics and drug dynamics in the organism are not included in the drug modelling, the simulated interventions produce an instant effect. Thus, lower waiting times are needed in-silico to understand whether a drug might be effective.

Moreover, as mentioned before, we simulated ideal pharmacological conditions, in which a complete SK and K_2_P channel inhibition was explored. While this might not be the case in clinical practice, we have ensured the APD and ERP prolongation in-silico match the results obtained in experimental and clinical conditions.

## Conclusion

Our prospective, large-scale in-silico trial study identifies key factors determining the success of combined versus single SK and K_2_P channel block, highlighting the polypharmacological inhibition as a safe and very effective strategy for AF management. Moreover, this study strengthens the power of in-silico trials based on human modelling and simulation for understanding optimal cardiac therapies.

